# Dampened PI3K/AKT signaling contributes to cancer resistance of the naked mole rat

**DOI:** 10.1101/2020.02.27.967729

**Authors:** Jing Zhao, Xiao Tian, Yabing Zhu, Zhihui Zhang, Elena Rydkina, Yongxuan Yuan, Hongyun Zhang, Bhaskar Roy, Adam Cornwell, Eviatar Nevo, Xiaoxiao Shang, Runyue Huang, Karsten Kristiansen, Andrei Seluanov, Xiaodong Fang, Vera Gorbunova

## Abstract

Mammalian species have a dramatically different susceptibility to cancer. However, how cancer-resistant species resist oncogenic transformation is not fully understood. Here, we performed a comprehensive analysis of oncogene-induced transcriptional changes in the fibroblasts of a cancer-prone species, the mouse, and three cancer-resistant species, the human, the blind mole rat, and the naked mole rat. We report that multiple cellular processes are more refractory to oncogene-induced transcriptional changes in blind mole-rat, naked mole-rat, or human cells compared to mouse cells, such as cell division, cell adhesion, extracellular matrix organization, and metabolism. Strikingly, naked mole rat cells are more resistant to Ras-induced transcriptional changes compared to the other three species. As a mechanism, we found that critical genes in the PI3K pathway including *Akt1* and *Pik3ca* are downregulated in naked mole rat cells. Activating the PI3K/AKT pathway in the naked mole rat cells renders them susceptible to tumorigenic transformation. This study provides multiple new insights into anti-cancer mechanisms in cancer-resistant species of mammals.

**Significance statement:** Animal species differ greatly in their cancer susceptibility. Cancer rates in the mouse range from 50-90%, while two other rodent species, the naked mole rat and the blind mole rat have only a few cancer cases ever reported. Here we examined the mechanisms responsible these differences by comparing changes in transcription patters in response to oncoproteins in the mouse, naked mole rat, blind mole rat and human cells. The most striking finding was that the naked mole rat cells were resistant to transcriptional changes induced by oncogenic Ras. We found that pathways downstream of Ras were naturally attenuated in the naked mole rat. This finding identifies a novel mechanism that evolved to provide tumor resistance to the naked mole rat.

## Introduction

Cancers are caused by mutations attributable to both intrinsic and extrinsic factors, such as erroneous DNA replication and environmental mutagens(1, 2). Mammalian species with larger body mass and longer lifespan have an increased risk of cancer because such species have a greater number of cells and more time to accumulate cancer-causing mutations. However, in reality, large and long-lived species have a lower incidence of cancer than small short-lived species, which is known as Peto’s paradox(3, 4). The solution for this paradox is that larger and longer-lived species have evolved additional anticancer mechanisms compared to small and short-lived species.

Cells from long-lived species require more oncogenic mutations to become malignantly transformed than cells from short-lived species(5). For example, mouse cells require only two oncogenic “hits”, which include inactivation of p53 or pRb tumor suppressor and expression of a constitutively active form of Ras oncogene, such as H-Ras V12(6, 7). In contrast, malignant transformation of human cells requires five “hits”, including inactivation of both p53 and pRb, expression of H-Ras V12 and telomerase, and constitutive activation of the PI3K pathway(8). This phenomenon is not restricted to mouse and human cells. In fact, by analyzing the requirements for malignant transformation in 18 rodent species, we recently showed that there is a general trend that more oncogenic “hits” are required to transform the cells from the longer-lived species(5). These results indicate that long-lived species harbor more anti-cancer mechanisms than short-lived species.

Multi-species comparative studies showed that some of these anti-cancer mechanisms were convergently evolved. For example, most of the large-bodied species (in general also longer-lived) have shorter telomeres and repress telomerase activity in their somatic cells(9, 10). As a result, telomeres shorten with each cell division and eventually cells undergo replicative senescence. This mechanism prevents unlimited cell division that is required for malignant transformation. Small long-lived species, however, have active telomerase in their somatic tissues. Instead of utilizing the replicative senescence mechanism, these species evolved unique telomere-independent anti-cancer mechanisms(4). For example, naked mole rats (NMR) (*Heterocephalus glaber*) and blind mole rats (BMR) (*Nannospalax galili*) are small long-lived rodent species that rarely develop cancers(11–14). To resist cancer, naked mole rat cells synthesize a large amount of high-molecular-weight hyaluronan, which could inhibit the oncogenic transformation induced by SV40 large T and H-Ras V12(15). It was characterized that the association with the hyaluronan receptor CD44 is critical for the tumor suppression effect conferred by high-molecular-weight hyaluronan. In contrast, blind mole rat cells utilize a unique interferon-mediated “concerted cell death” mechanism to prevent hyperplasia(14).

The diversity of anti-cancer mechanisms in different long-lived species prompted us to perform a multi-species comparison of the transcriptional network in response to oncogenic insults. This allowed us to examine both the common and species-specific transcriptional changes at different stages of oncogenic transformation. In this study, we analyzed the transcriptome of the mouse, human, naked mole-rat, and blind mole-rat cells during the multi-step oncogenic transformation. We report that multiple cellular and molecular processes are more resistant to changes stimulated by oncogenic insults in blind mole-rat, naked mole-rat, or human cells, including cell division, cell adhesion, extracellular matrix organization, and metabolic process etc. Specifically, we found that critical genes in the PI3K pathway, such as *Akt1* and *Pik3ca*, are downregulated in the naked mole-rat cells, which contributes to their resistance to Ras-induced transcriptional changes. This study provides multiple new insights into the anti-cancer mechanisms in cancer-resistant species.

## Results

### Experimental setup for RNA-seq analysis of oncogene-induced gene expression changes

To obtain a comprehensive picture of gene expression changes upon oncogenic insults, we performed transcriptomic analysis on the mouse, human, naked mole-rat, and blind mole-rat cells at each step of a multi-step oncogenic transformation. Skin fibroblasts from mice, naked mole rats and blind mole rats were transformed with SV40 large T (LT) alone, or together with H-Ras V12. Fibroblasts from these species express active telomerase, while human cells do not, and require reactivation of telomerase expression for malignant transformation(8). Without telomerase, human cells transformed with LT and oncogenic Ras undergo crisis(5). Therefore, human fibroblasts HCA2 were first immortalized by overexpressing human telomerase (hTERT), and then transformed with LT alone, or together with H-Ras V12. The tumorigenicity of the resulting cell lines has been tested previously in mouse xenograft models(5) and is summarized in Table 1. Briefly, LT and H-Ras V12-transformed mouse (M-L-R) and blind mole-rat skin fibroblasts (BMR-L-R) form tumors in immunodeficient mice, whereas LT and H-Ras V12-transformed naked mole-rat skin fibroblasts (NMR-L-R) do not robustly form tumors(15, 16) unless high-molecular-mass hyaluronan (HMM-HA) is abolished(15). hTERT, LT and H-Ras V12-transformed human skin fibroblasts (H-T-L-R) do not form tumors.

**Table 1.**
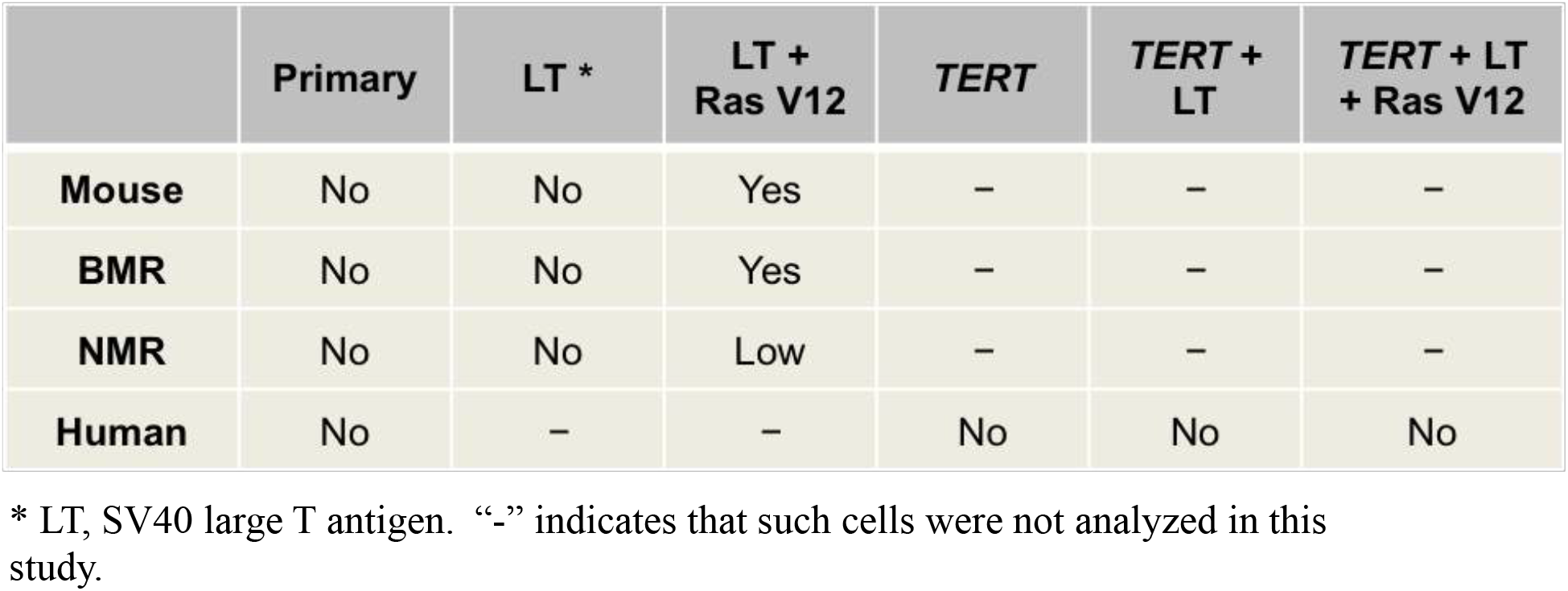
Tumorigenicity of the cells used for RNA sequencing

The list of cell lines subjected to RNA sequencing is shown in Table S1. Three replicates of each cell line were used for RNA sequencing. All libraries were sequenced on Illumina HiSeq platform. After filtering adapters and low-quality reads, a total number of 729,359,058 pairs of reads were obtained. Clean reads of all samples of each species were aligned against corresponding reference genomes using HISAT2, and gene expression levels were calculated using Cufflinks. In total, expression of 15426, 16073, 15833 and 15249 genes were detected in mouse, human, naked mole rat and blind mole rat cells, respectively. The number of genes detected in each sample is indicated in Table S1. The replicates for each treatment within each species showed high Pearson correlation coefficients ranging from 0.967 to 0.999 (Fig. S1).

In order to compare the RNA-seq data across species, we identified orthologs across all four species using mouse genes as the reference. First, we aligned protein sequences from each species against the mouse protein sequences. Next, we identified orthologs between mouse and each species according to gene identities. Finally, we merged the three outputs into one final table. In total, 13276 orthologs were shared by all four species (Table S2).

### Gene expression is significantly influenced by oncogenic insults

To assess the expression changes induced by oncogenes, we performed Principal Component Analysis (PCA) across all samples of the same species. Gene expression in transformed cells clearly segregated from primary cells in all four species, indicating that transforming factors, LT and H-Ras V12, significantly influence gene expression (Fig. 1). However, the segregation patterns in the four species were different. Specifically, NMR and human cells, which do not efficiently form tumors after transformation with LT and H-Ras V12, had different segregation patterns compared to mouse and BMR cells. While mouse and BMR cells underwent changes with each transforming step, human cells only exhibited major expression changes upon introduction of three oncogenic “hits” (TLR). In contrast, for NMR cells, LT (L) and LT+Ras (LR) transformed cells grouped together, indicating that LT, but not Ras, is responsible for the major expression differences between naked mole rat samples. This suggests that NMR cells are refractory to the expression changes induced by Ras.

**Figure 1.**
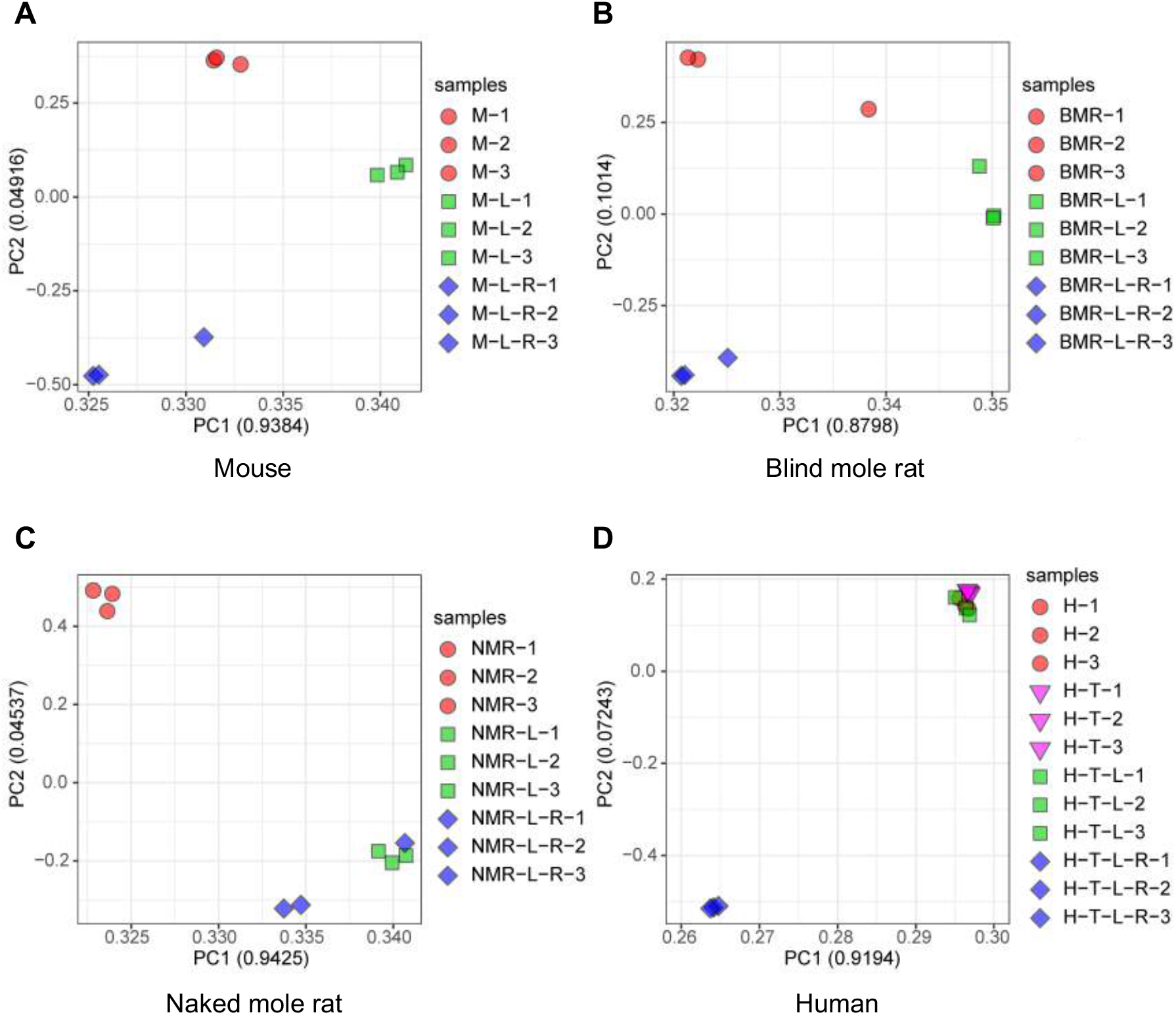
Principal component analysis (PCA) of the RNA-seq datasets from the four species. PCA was done using the expression of 13276 orthologous genes across all four species. The first two Principal Components of each analysis were extracted. Values in parenthesis indicate the variance explained by each of the PCs. M, mouse; BMR, blind mole rat; NMR, naked mole rat; H, human; L, SV40 LT antigen; R, H-Ras V12; T, telomerase.

### Cell cycle-related genes show differential response to oncogenes between species

The transforming mechanisms of LT and oncogenic Ras have been extensively characterized(8, 17). LT binds and inactivates p53 and the Rb family of tumor suppressors. Ras is a frequently mutated proto-oncogene in human tumors. The constitutively active Ras protein triggers multiple downstream signaling cascades through its interaction with three major Ras effector proteins, Raf, PI3K, and RalGEFs(18). Considering that the LT and H-Ras V12 co-expressing cells of the four species have different tumorigenic capacity (Table 1), we set out to compare their gene expression across the four species, aiming to identify genes or pathways that prevent tumorigenesis in NMR and human cells compared to mouse and BMR cells.

The average expression value of each gene from three replicates of each treatment was used to determine differential expressed genes (DEGs). More than 2,000 genes showed differential expression in every pairwise comparison among the LT and Ras-transformed cells between species (Table S3). Next, we performed enrichment analysis on the DEGs revealed within each pair. We found that multiple gene ontology (GO) terms including cell cycle, cell division, and mitotic nuclear division were significantly down-regulated in non-tumorigenic NMR-L-R cells compared to tumorigenic M-L-R or BMR-L-R cells (Fig. 2A, Fig. S2A). Furthermore, the down-regulated genes in the “cell cycle” term were involved in the regulation of different cell cycle stages (Fig. 2B), suggesting that multiple molecular mechanisms ensure that naked mole-rat cells expressing LT and Ras do not accelerate their cell cycle. Evading growth suppression is a hallmark of cancer(19). The failure to fully activate cell cycle-related genes by LT and oncogenic Ras reflects that there are intrinsic mechanisms in naked mole-rat cells that make them resist this oncogene-induced transcriptional change. Interestingly, H-T-L-R cells, which are also non-tumorigenic, showed similar downregulation in cell cycle-related genes compared to M-L-R cells (Fig. 2C). Such genes were not among the most significantly enriched GO terms when NMR-L-R cells were compared with H-T-L-R cells (Fig. S2B).

**Figure 2.**
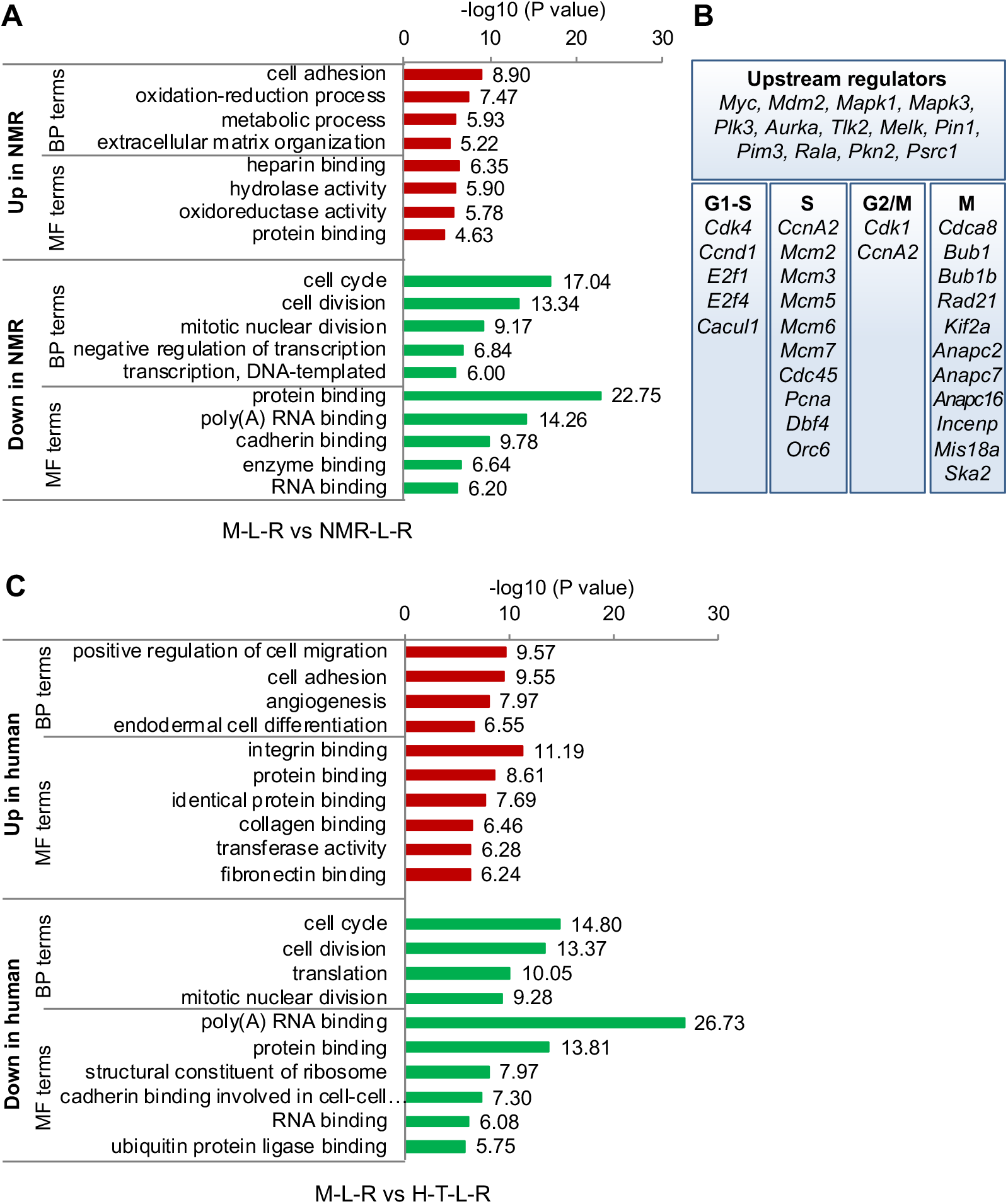
Gene expression variation of the SV40 Large T and Ras-transformed cells across species. (**A**) GO enrichment analysis between mouse and naked mole rat transformed cells expressing Large T and H-Ras V12 (M-L-R vs NMR-L-R). Biological Process (BP) and Molecular Function (MF) terms with the most significant P values were plotted. (**B**) The downregulated “cell cycle” genes in the NMR-L-R cells (derived from **A**) were categorized into cell cycle stages based on when their functions are executed. (**C**) GO enrichment analysis between mouse and human transformed cells with Large T and H-Ras V12 (M-L-R vs H-T-L-R). Biological Process (BP) and Molecular Function (MF) terms with the most significant P values were plotted.

Surprisingly, though both M-L-R and BMR-L-R cells are tumorigenic (Table 1), cell cycle-related GO terms, such as cell cycle, cell division, regulation of cell cycle, and mitotic nuclear division were significantly down-regulated in BMR-L-R cells compared to M-L-R cells (Fig. S2C), suggesting that BMR-L-R cells are less malignant than M-L-R cells. Indeed, our previous publication showed that BMR-L-R cells take a longer time than M-L-R cells to form a similar-sized tumor in immunodeficient mice(5), indicating that BMR transformed cells are less malignant than mouse transformed cells.

### Cell adhesion- and metabolic process-related genes are upregulated in naked mole rat cells

Upregulated genes in NMR-L-R cells relative to M-L-R cells were enriched for multiple GO terms, among which the most significant was “cell adhesion” (Fig. 2A). Interestingly, this group of upregulated genes in NMR includes *Has2, Hapln1, Lamc1, Col7a1, Col6a2, Col6a1, Col8a1,* and *Itga2b*, which are responsible for synthesizing or regulating extracellular matrix components. This result supports that the extracellular matrix of naked mole rat cells plays an important role in preventing migration and invasion that occur during malignant transformation. In line with this notion, we previously found that knock-down of HA synthesis or enzymatic degradation of HA, which destroy the scaffold structure of the extracellular matrix, is sufficient to enable the tumor formation by the NMR-L-R cells(15). Other upregulated genes in NMR-L-R cells were enriched in pathways including “oxidation-reduction process” and “metabolic process” (Fig. 2A). Since oncogenic Ras and the Ras effector pathways contribute to the metabolic reprograming towards anabolic processes(20, 21), the relative upregulation of oxidation-reduction genes in NMR-L-R compared to M-L-R suggest that cancer resistance of naked mole-rat cells may also be at least partially due to the resistance of NMR cells to metabolic transformation.

### Oncogenic Ras induces fewer differential expressed genes in naked mole-rat cells

Next, we examined gene expression changes triggered by LT and Ras within each species. The transcriptome profiles of LT and H-Ras V12 co-expressing cells were compared to the corresponding primary cells. The genes with differential expression changes of more than 2-fold with *p* values < 0.01 were extracted, which represents a strict threshold to reduce the false positive identification of DEGs. The number of LT and Ras-induced DEGs was similar in the mouse, blind mole-rat, and human cells, but was much lower in naked mole-rat cells (Fig. 3A).

**Figure 3.**
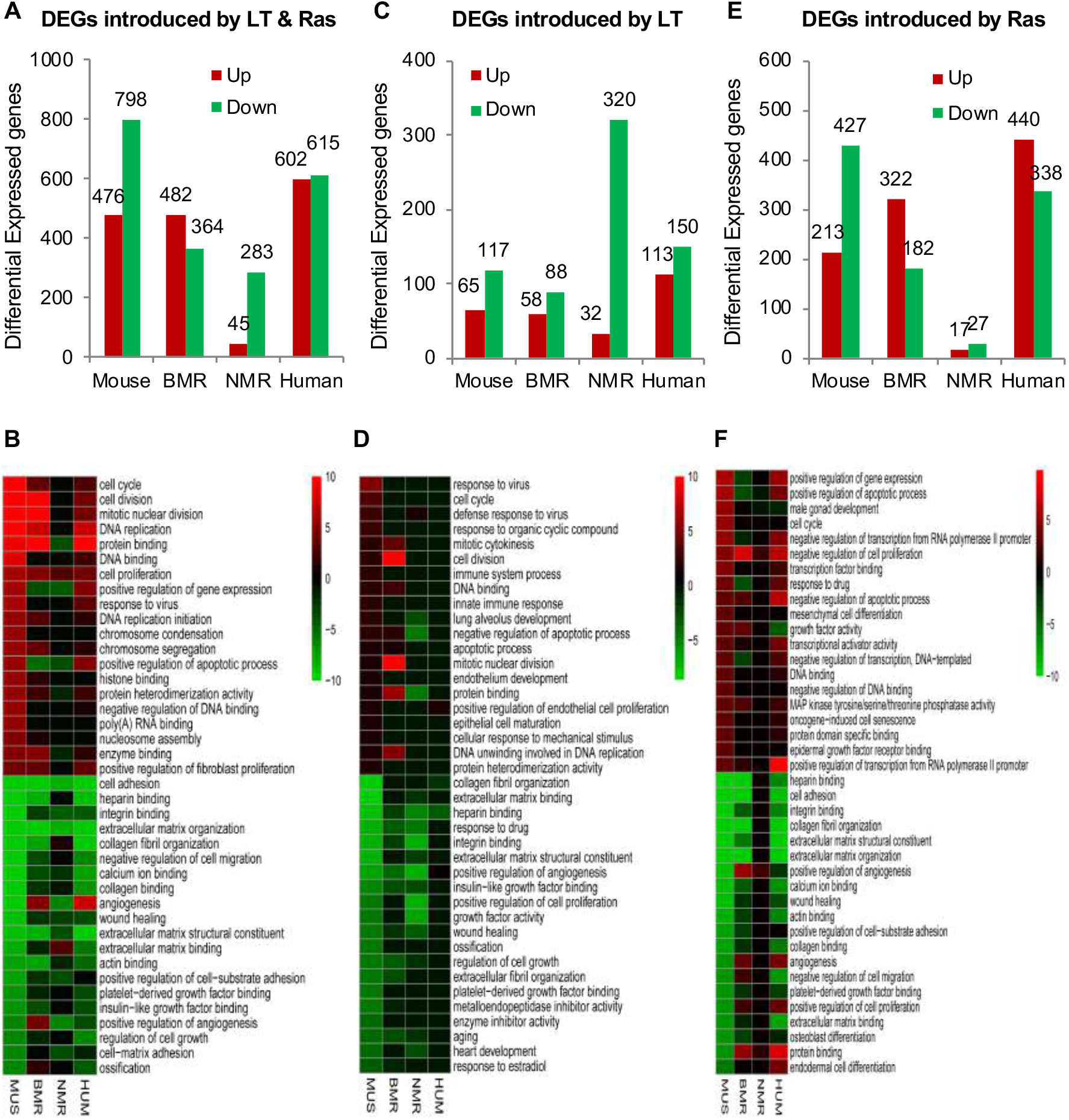
Differentially expressed genes induced by oncogenes across species. (**A**) The number of differentially expressed genes (DEGs) induced by the combination of SV40 Large T (LT) and H-Ras V12 (Ras) across species. (**B**) GO enrichment analysis of the DEGs derived from **A**. The GO terms are arranged in the order of decreased significance in the mouse cells. The top 20 significantly upregulated (red) and downregulated (green) terms are shown. The Biological Process (BP) and Molecular Function (MF) terms are included. (**C**) The number of differentially expressed genes induced by LT across species. (**D**) GO enrichment analysis of the DEGs derived from **C**. (**E**) the number of differentially expressed genes induced by H-RasV12 across species. (**F**) GO enrichment analysis of the DEGs derived from **E.** The GO terms were arranged in the order of decreased significance in the mouse cells.

Next, we performed enrichment analyses on LT and Ras-induced DEGs. In mouse cells, the LT and Ras-induced DEGs were enriched in multiple terms including upregulated cell cycle, cell division, and mitotic nuclear division, and downregulated cell adhesion, heparin binding, and integrin binding (Fig. 3B). These transcriptional changes are highly consistent with the known functions of p53, pRb, and H-Ras V12. However, in naked mole-rat cells, multiple corresponding terms were not significantly altered or altered to a much lesser extent (Fig. 3B), especially in the case of upregulated genes. Arranging the terms by the significance within NMR sample comparisons also indicated that NMR cells have a unique response to LT and Ras transformation (Fig. S3A).

We next dissected the transforming effect of LT and H-Ras V12 separately. The LT-transformed cells were compared with the corresponding primary cells, while the LT and H-Ras V12 co-expressing cells were compared with the LT-transformed cells. Interestingly, LT induced more down-regulated DEGs in the naked mole-rat cells than in the other three species (Fig. 3C). Enrichment analysis showed that the LT-induced down-regulated genes were more significantly enriched and grouped into different functional terms than up-regulated genes in all four species (Fig. 3D). Interestingly, the most significant LT-induced down-regulated terms in NMR, including “extracellular matrix organization”, “angiogenesis”, and “collagen catabolic process”, were unique compared to the other three species (Fig. S3B).

H-Ras V12 induced a much lower number of DEGs in the NMR cells (Fig. 3E). H-Ras V12-induced DEGs were enriched for multiple terms in the mouse, blind mole-rat, and human cells, including upregulated “cell cycle”, “growth factor activity”, etc. and downregulated “cell adhesion”, “ECM organization”, etc. (Fig. 3F). However, in the naked mole-rat cells, such terms were not significantly altered or altered to a much smaller extent (Fig. 3F and S3C). These results revealed that NMR cells display unique responses to both LT and oncogenic Ras: LT causes downregulation of more genes compared to the other three species, while oncogenic Ras fails to trigger massive expression changes.

### Critical genes in the PI3K pathway are downregulated in naked mole-rat cells

Since we found that NMR cells are refractory to oncogenic Ras, and we used human H-Ras for all the species, we tested whether differences in the H-Ras sequence between species are responsible for this lack of response. The alignment of H-Ras protein sequences between the species revealed that H-Ras proteins were identical between the mouse, blind mole rat, and human, whereas the naked mole rat H-Ras had Serine (instead of Glycine) at amino acid (aa) 138 (Fig. 4A).

**Figure 4.**
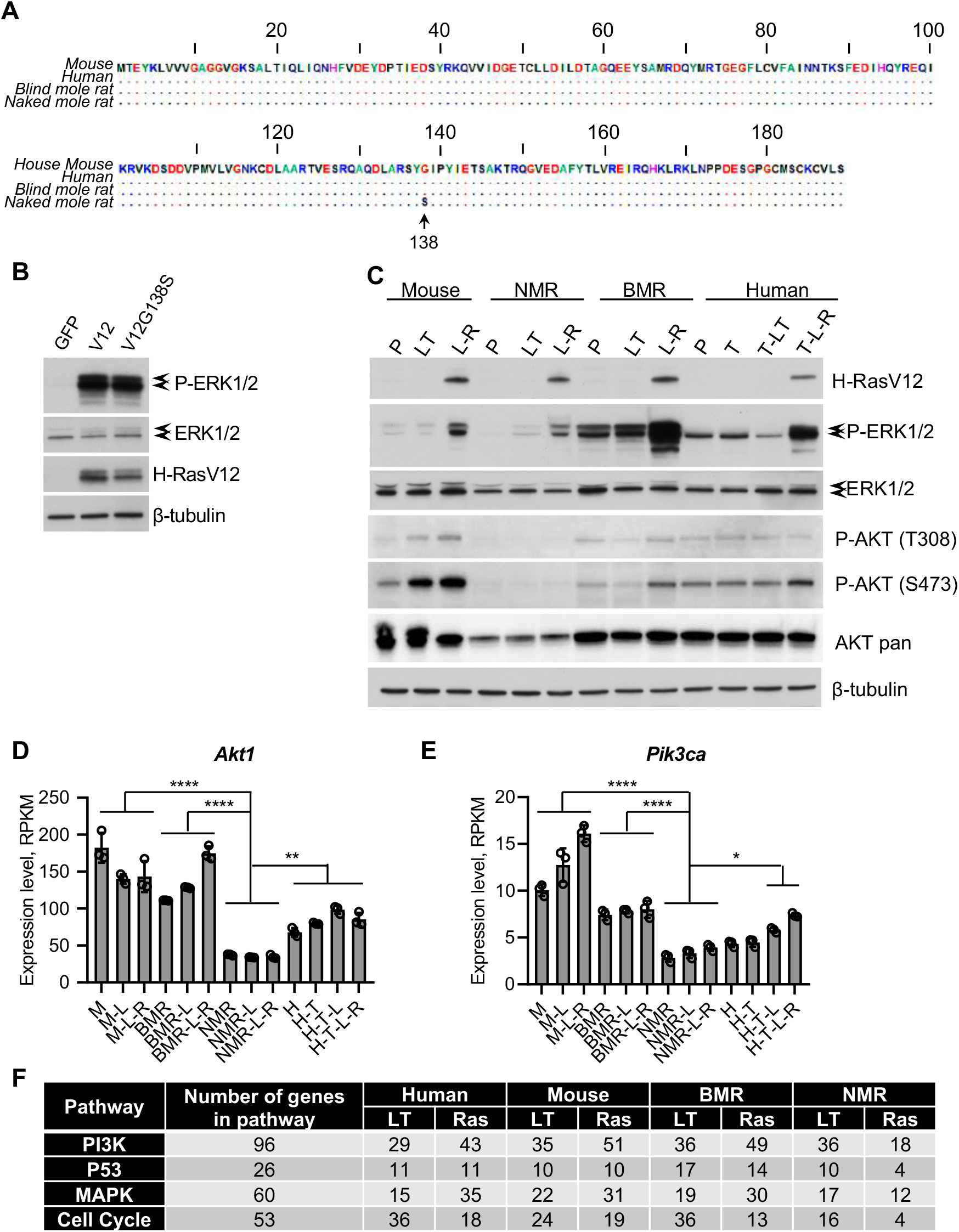
The PI3K/AKT pathway is downregulated in naked mole rat cells. (**A**) Alignment of the H-Ras protein sequences from mice, humans, blind mole rats, and naked mole rats. (**B**) Transient overexpression of both human (V12) and naked mole rat (V12G138S) H-Ras V12 proteins stimulate the ERK pathway. Naked mole rat cells were transiently transfected with 5 μg of indicated genes. Two days after transfection, cell lysates were used for Western blot. (**C**) The PI3K/AKT pathway is downregulated in naked mole rat cells. The stable cell lines (the same cells that were used for RNA-seq) expressing indicated oncogenes were used for Western blot. P, primary cells; LT, SV40 LT transformed; L-R, SV40 LT and H-Ras V12-expressing cells; T, telomerase; T-LT, telomerase and SV40 LT expressing cells; T-L-R, telomerase, SV40 LT and H-Ras V12 expressing cells. (**D**) The transcription of *Akt1* is downregulated in naked mole rat cells. The RPKM value of *Akt1* from each sample was extracted from the RNA-seq data. Statistical significance was determined using one-way ANOVA with Tukey’s post hoc test. The least significance values among each group of comparison were indicated. **, p < 0.01; ****, p < 0.0001. (**E**) The transcription of *Pik3ca* is downregulated in naked mole rat cells. Statistical significance was determined using one-way ANOVA with Tukey’s post hoc test. The least significance values among each group of comparison were indicated. *, p < 0.05; ****, p < 0.0001. (**F**) The number of DEGs induced by SV40 LT or H-Ras V12 in each species.

To test whether the difference at aa 138 is responsible for the lack of activation of the Ras pathway in the NMR cells, we introduced the G138S mutation into human H-Ras V12, generating V12G138S mutant. This H-Ras mutant encodes the same protein as the NMR H-RasV12 oncogene. We next compared the ability of human and NMR versions of oncogenic RAS to trigger phosphorylation of ERK, the downstream effector of Ras, in the NMR cells. Remarkably, the two versions of H-Ras similarly stimulated phosphorylation of ERK when overexpressed in the NMR cells (Fig. 4B). This result indicates that human H-Ras V12, which was used to generate the NMR-L-R cells, is functional in the NMR cells.

Ras has multiple downstream effectors. The Raf serine/threonine kinases are a family of the best characterized Ras effector proteins. Activated Raf phosphorylates and activates MEK1/2 dual-specificity kinases, which then activate the ERK1/2 mitogen-activated protein kinases (MAPKs)(22). We characterized the activity of the MAPK pathway in the cells that were used for RNA sequencing by testing the phosphorylation levels of the ERK proteins. We found that the phosphorylation level of ERK1/2 is similar between NMR-L-R and M-L-R cells (Fig. 4C), indicating that the ERK pathway is activated by Ras in NMR cells. Interestingly, the level of phosphorylated ERK is much higher in BMR cells compared to the other three species (Fig. 4C). The high activity of the ERK pathway in the primary BMR cells may contribute to the fast cell growth before the concerted cell death (CCD) that we previously reported (14).

The phosphoinositide 3-phosphate lipid kinase (PI3K) is another major effector protein of Ras(23). PI3K catalyzes the conversion of phosphatidylinositol 4,5-phosphate (PIP2) to phosphatidylinositol 3,4,5-phosphate (PIP3). Interaction of AKT with PIP3 facilitates the phosphorylation and activation of AKT by PDK1 and mTOR complex 2 (mTORC2) (24). Activated AKT promotes cell cycle progression and cell survival by phosphorylating a long list of protein targets(24). In human cells, constitutively active PI3K/AKT pathway is required for tumorigenic transformation(8). We found that the phosphorylation levels of AKT on T308 and S473 were much lower in naked mole-rat cells compared to the cells of the other three species (Fig. 4C), indicating a suppressed PI3K pathway in naked mole rat cells. Importantly, H-Ras V12 increased the phosphorylation level of AKT at S473 in mouse, blind mole rat, and human cells, but not in naked mole rat cells (Fig 4C). Furthermore, we found that the total protein level of AKT was much lower in NMR samples (Fig. 4C). The lower transcription of *Akt1* in NMR samples was also revealed in our RNA-seq data (Fig. 4D). In addition, an important PI3K gene, *Pik3ca,* which encodes a catalytic subunit of PI3K, showed significant down-regulation in naked mole-rat cells (Fig. 4E).

Considering the pivotal roles of PI3K and AKT in the PI3K pathway, we suspected that the downregulation of such genes may resist H-Ras V12-induced activation of the PI3K pathway. Indeed, though LT induced a similar number of DEGs in the PI3K pathway across all four species, H-Ras V12 induced much fewer DEGs among the PI3K/AKT targets in NMR cells (Fig. 4F and S4).

### Activating the PI3K/AKT pathway promotes tumorigenicity in naked mole rat cells

The strong suppression of the AKT pathway in NMR cells suggested that it may contribute to the low level of tumorigenicity induced by LT and oncogenic Ras. We next tested whether activating the AKT pathway in NMR cells could promote tumorigenic transformation. SV40 small T (ST) antigen binds and inactivates protein phosphatase 2A (PP2A), thereby activating the PI3K/AKT pathway(25). We generated stable cell lines expressing ST under the control of a weak promoter (retroviral LTR promoter) or a stronger CAG promoter. Interestingly, expressing a high level of ST under the CAG promoter in NMR-L-R cells increased the tumorigenicity of NMR-L-R cells from ~ 29% (4 tumors formed out of 14 xenografts) to ~75% (9 tumors formed out of 12 xenografts). However, the ST driven by the weaker promoter, LTR, failed to enhance the tumorigenicity of NMR-L-R cells (Fig. 5A). Furthermore, NMR cells expressing LT and ST failed to form tumors (Fig. 5A), indicating that ST cannot substitute oncogenic Ras in NMR cell transformation. The combination of LT, ST, *TERT*, and H-Ras V12 driven by the CAG promoter sufficiently transformed human foreskin fibroblasts (Fig. 5A). Interestingly, though high expression of ST in NMR (LT+RAS)^LTR^ cells significantly improved tumorigenicity, tumors formed from both NMR (LT+RAS)^LTR^ and NMR (LT+RAS)^LTR^+ST^CAG^ cell lines have a very long latency (~5 weeks) compared to fully-transformed mouse and human cells (Fig. 5B). Both the low frequency and long latency of tumor formation from the NMR (LT+RAS)^LTR^ cells suggest that additional mutations may accumulate in a small fraction of the population during cell passage that overcome the hyaluronan barrier and other potential tumor suppressing mechanisms.

**Figure 5.**
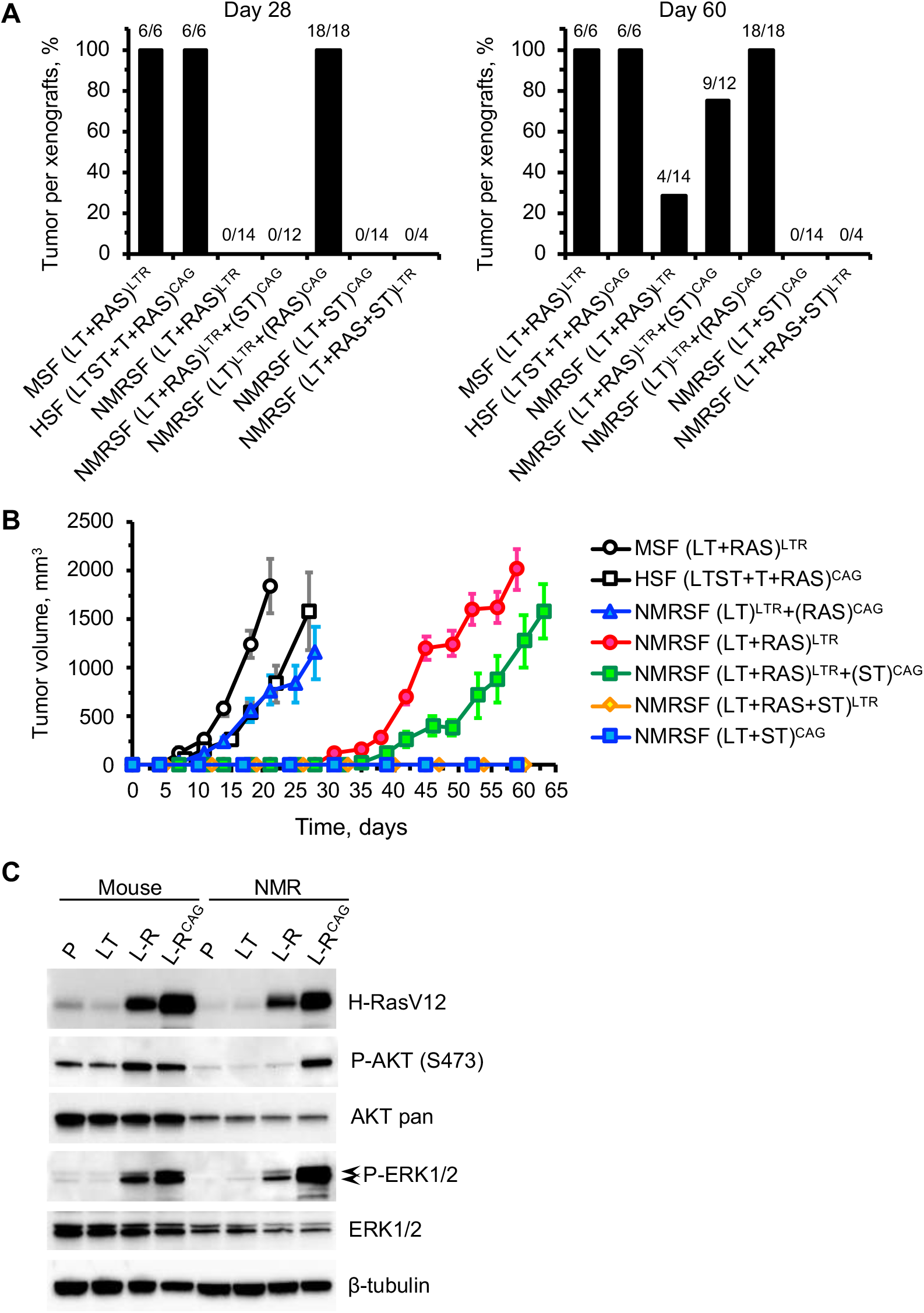
Activating PI3K/AKT pathway promotes tumorigenicity of the naked mole rat cells. (**A**) Frequency of tumor formation of xenografts of mouse, human, and naked mole rat cells expressing different combinations of oncoproteins and either low or high expression of H-Ras V12 and SV40 Small T antigen. LTR a is a weak retroviral promoter that is widely used for malignant transformation of mouse and human cells. CAG is a strong synthetic promoter. The numbers above the bars indicate number of tumors formed/number of injection sites detectable by day 28 and by day 60 post injection. (**B)** Growth curves of tumors formed by the transformed cells in **A**. The error bars indicate SEM. (**C**) Western blot analysis of AKT and ERK phosphorylation in mouse and NMR cells expressing different levels of H-Ras V12. P, primary cells; LT, SV40 LT transformed; L-R, cells expressing SV40 LT and H-Ras V12 under LTR promoter. L-R^CAG^, cells expressing SV40 LT and H-Ras V12 under CAG promoter.

PI3K is a major effector protein of the Ras oncogene(7). It was previously found that a higher expression level of H-Ras V12 promotes human cell transformation(26). We next asked if the PI3K/AKT pathway could be activated by a higher expression level of oncogenic Ras in NMR cells. H-Ras V12 driven by a CAG promoter was stably introduced into NMR-L cells, generating NMR-L-R^CAG^ cells. The resulting cell lines expressed a higher level of H-Ras V12 than the NMR-L-R cells where LT and H-Ras V12 were driven by retroviral LTR promoters (Fig. 5C). As a result, the AKT pathway is strongly activated gauged by the phosphorylation level of AKT at S473 (Fig. 5C). NMR-L-R^CAG^ cells formed tumors 100% of the time (18 tumors formed out of 18 xenografts) and displayed significantly reduced latency (Fig. 5A and 5B). Tumors formed from each injection are summarized in Fig. S5.

Both lines of results, increased tumorigenicity of NMR cells expressing LT+RAS+ST^CAG^ and NMR cells expressing LT+Ras^CAG^, demonstrate that forced activation of the AKT pathway overrides the resistance of NMR cells to malignant transformation. Therefore, we conclude that the naturally suppressed AKT pathway in NMR cells contributes to their resistance to Ras-induced tumorigenic transformation.

## Discussion

Animal species differ dramatically in lifespan and cancer susceptibility(27). Long-lived species delay the onset of cancer and are expected to have more advanced tumor suppressor mechanisms. In line with this notion, we and others have shown that cells from long-lived species are more resistant to oncogenic transformation than that from short-lived species(5, 7, 8, 28). However, the molecular mechanisms underlying cancer resistance in long-lived species are still largely a mystery.

In this study, we analyzed transcriptional changes that occur at different stages of malignant transformation in mouse, human, blind mole rat and naked mole rat cells. The four species showed remarkably different dynamics of the transcriptome changes. Mouse cells drastically changed expression patterns in response to both LT and oncogenic Ras. Blind mole rat cells were more similar to mouse cells in their responses. Human cells were refractory to changes induced by LT, but responded strongly to Ras, while naked mole rat cells changed expression pattern in response to LT but were strikingly resistant to oncogenic Ras.

Transcriptional changes that occurred in response to oncogenic Ras included genes regulating extracellular matrix organization, cell cycle, and metabolism. Extracellular matrix remodeling promotes mitotic growth and activates invasion and metastasis signaling in transformed cells(19). Our results suggest that maintaining homeostasis of the extracellular matrix is a critical mechanism to resist tumor formation. Interestingly, naked mole-rat cells produce a unique extremely high-molecular-weight hyaluronan molecule. Degrading this molecule promotes tumor formation in naked mole-rat cells(15).

The most striking difference between the species was the resistance of the naked mole rat cells to oncogenic Ras. Remarkably, further transcriptome analysis showed that naked mole-rat cells significantly suppress the AKT pathway by inhibiting the expression of multiple pivotal mediators in the PI3K/AKT signaling cascade, including *Akt1* and *Pik3ca*. Previous studies have demonstrated the requirement of activating the PI3K/AKT pathway in transforming human cells(7, 8, 25, 29). Therefore, attenuated PI3K/AKT signaling cascade in the naked mole rat provides an explanation for the resistance of naked mole-rat cells to Ras-induced transcriptional changes, as PI3K is a major downstream effector of Ras.

Naked mole rat fibroblasts are not readily transformed by a combination of LT and oncogenic Ras(15). However, here we found that stimulating the AKT pathway by SV40 ST or by expressing a very high level of oncogenic Ras significantly promotes the oncogenesis of naked mole-rat cells. Similar observations were previously reported for human cells, where tumorigenicity of the primary cells transformed with LT and oncogenic Ras was dependent on the level of expression of Ras oncogene(26).

Our results, for the first time, establish a critical role of the dampened PI3K/AKT pathway in mediating cancer resistance in naked mole rat cells. The PI3K/AKT signaling pathway has also been recognized to regulate longevity. In *C. elegans* and Drosophila, abolishing PI3K/AKT signaling significantly extends lifespan(30, 31). In mice, partial inactivation of PI3K or AKT enhances metabolic function and extends lifespan(32, 33). Considering the tight connection to lifespan regulation, the natural suppression of the PI3K/AKT pathway in the naked mole rat is likely an adaption to achieve not only cancer resistance, but also a longer lifespan.

## Methods

### Animals

C57BL/6 mice were purchased from the Jackson Laboratory. The naked mole rats and the blind mole rats were from the University of Rochester colonies. NIH-III nude mice (Crl:NIH-*Lyst*^*bg-J*^*Foxn1*^*nu*^*Btk*^*xid*^) were purchased from Charles River Laboratories. All animal experiments were approved and performed in accordance with the guidelines set up by the University of Rochester Committee on Animal Resources.

### Cell extraction and culture

Primary fibroblasts from mice, naked mole rats, and blind mole rats were isolated from underarm skin samples. The mouse samples were from a C57BL/6 mouse. The naked mole-rat and the blind mole-rat skin samples were from the University of Rochester colonies. The human skin fibroblasts were a gift from Pereira-Smith lab at University of Texas Health Science Center at San Antonio. Mouse, human, and blind mole-rat cells were cultured at 37°C with 5% CO_2_ and 3% O_2_. Naked mole rat cells were cultured at 32 °C (body temperature of naked mole rats) with 5% CO_2_ and 3% O_2_. All cells were cultured in EMEM medium (ATCC) supplemented with 15% (vol/vol) fetal bovine serum (Gibco) and 1× penicillin-streptomycin (Corning).

### Transfections

Cells were plated at 5×10^5^ cells per 10 cm plate 2 days before transfection, except for naked mole rat cells which were seeded at 2×10^5^ cells per plate 5 days before transfection. Cells were harvested, resuspended in NHDF transfection solution (Amaxa) and transfected with corresponding plasmid DNA using Amaxa Nucleofector II on program U-20.

### Plasmids and stable cell line generation

Mouse and naked mole-rat primary and transformed cells used for RNA sequencing were previously published(15). Basically, Not1-linearized pBabe-puro largeTcDNA plasmid (Addgene plasmid # 14088) was transfected into mouse and naked mole-rat skin fibroblasts and selected with puromycin for 2-4 weeks (mouse cells took ~2 weeks, and naked mole-rat cells took ~4 weeks) to generate stable clones. After selection, stable clones were pooled to minimize the clonal variations. Pooled clones were expanded before being transfected with Not1-linearized pWZL-hygro H-Ras V12 plasmid (Addgene plasmid # 18749) and selected with hygromycin for 2-4 weeks to generate stable clone. The expression of each protein was confirmed by western blotting. Blind mole-rat cells used for RNA sequencing were generated using the same method. Human cells used for RNA sequencing were first integrated with Not1-linearized pBABE-neo-hTERT plasmid (Addgene plasmid # 1774) and selected with G418. The generated HCA2-hTERT cells were integrated with LT and H-Ras V12 using the same method described above. For each antibiotic selection, a non-transfected control was included to ensure complete death of the un-transfected cells. The oncogenes were all driven by retroviral LTR promoter.

To generate stable cell lines for xenograft experiments, two different techniques were used. The LTR-driven plasmids expressing LT (Addgene plasmid # 14088), H-Ras V12 (Addgene plasmid # 18749), and ST (Addgene plasmid # 8583) were integrated into cells using the same transfection and selection method described above. Not1-linearized pBABE-zeo small T integration was selected using Zeocin. The CAG-driven oncogenes were integrated into the cells using piggyBac (pPB) expression vectors described previously(5). Basically, to generate HSF (LTST+T+RAS)^CAG^ cells, HCA2 was first co-transfected with pPB-puro-LTST and piggyBac transposase (PBase) plasmids and selected with puromycin. Stable clones were pooled and co-transfected with pPB-neo-hTERT and PBase plasmids and selected with G418. The generated stable clones were pooled and then co-transfected with pPB-hyg-H RasV12 and PBase plasmids and selected with hygromycin. To generate NMRSF (LT)^LTR^+(RAS)^CAG^ cells, the NMRSF (LT)^LTR^ cells generated using pBABE plasmid (Addgene plasmid # 14088) above were co-transfected with pPB-hyg-H RasV12 and PBase plasmids and selected with hygromycin. To generate NMRSF (LT+RAS)^LTR^+(ST)^CAG^ cells, NMRSF (LT+RAS)^LTR^ described above was co-transfected with pPB-zeo-ST and PBase plasmids and selected with Zeocin. To generate NMRSF (LT+ST)^CAG^ cells, NMRSF was first co-transfected with pPB-puro-LT and PBase, and selected with puromycin. The generated stable clones were co-transfected with pPB-zeo-ST and PBase and selected with Zeocin. Extensive passages were avoided to prevent clonal expansion.

### Xenograft assay and tumor measurement

Xenograft experiments were performed according to our previously published procedure(15) with minor modifications. Basically, two- to three-month-old female NIH-III nude mice (Crl:NIH-*Lyst*^*bg-J*^*Foxn1*^*nu*^*Btk*^*xid*^) were used to establish xenografts. Cells were re-suspended to a dilution of 2 × 10^6^ cells in 100 μl ice-cold 50% matrigel (Corning) in PBS. 100 μl of cell suspension was injected subcutaneously into the flank using a 22-gauge needle after general anesthesia. The mice were monitored and the size of tumors was measured using a caliper twice a week. When the length of the tumors reached 20 mm, the mice were euthanized. If no tumors grew, the mice were euthanized after two months. The volume of the tumors was calculated by the following formula: Tumor volume (mm^3^) = D × d × d /2, where D and d are the longest and shortest diameter of the tumors in mm.

### Antibodies

The following antibodies were used in this study: phospho-ERK1/2 (4370, Cell Signaling Technology (CST)), ERK1/2 (4695, CST), phospho-T308-AKT (13038, CST), phospho-S473-AKT (4060, CST), AKT1 (2938, CST), AKT (4691, CST), H-Ras V12 (ab140962, Abcam), β-tubulin (ab6046, Abcam).

### RNA extraction and sequencing

Total RNA was extracted from cultured cells using Qiagen RNeasy mini kit and treated with DNase I (DNA-free, Ambion). Oligo (dT) was used to isolate mRNA. All samples were mixed with the fragmentation buffer. The mRNAs were fragmented and 1 μg was used for cDNA synthesis (iScript, Bio-Red) in 20 μl reaction using the mRNA fragments as templates. Short fragments were purified and dissolved with EB buffer for end preparation and single nucleotide A (adenine) addition. After that, the short fragments were connected with adapters. The suitable fragments are selected for the PCR amplification. During the quality assessment steps, Agilent 2100 Bioanaylzer and ABI StepOnePlus Real-Time PCR System (Applied Biosystems) were used in quantification and qualification of the sample libraries. The RNAseq libraries were prepared with Illumina’s TruSeq RNA sample Prep kit according to the manufacturer’s protocols. Next, the libraries were taken for sequencing using Illumina HiSeq 4000.

### Data filtering and processing

After acquiring the sequencing data, Paired End reads (PE) were filtered to trim adapters and low-quality reads using SOAPnuke V1.5. We obtained a total number of 729,359,058 pairs of clean reads. After that, Human, mouse, NMR, and BMR RNA-Seq reads were first mapped to their respective reference genomes using HISAT2 and calculated FPKM values with Cufflinks. In total, 15426, 16073, 15833, and 15249 genes were expressed in mouse, human, naked mole-rat and blind mole-rat, respectively.

### Orthologs identification

To identify human, mouse, BMR, and NMR orthologous genes, we compared the RNA-seq data between species, and used mouse genes as the reference. First, we aligned mouse gene sequences against genes of other three species. Next, we identified orthologs between mouse and each species according to gene identities. Finally, we merged the three outputs into one final table. In total, 13276 orthologs were shared by all four species.

### Gene differential expression analysis

We conducted a cross-species differential expression gene analysis (DEG) using DESeq2, which takes replicates as one group. The thresholds for significance are fold change ≥ 2 and adjusted p-value ≤ 0.01. We did comparisons of L_R VS P, L VS P, and L_R VS L for each species, in which P means primary cells; L means LT-expressing cells; and L_R means LT and Ras-expressing cells. We also compared same processed samples between each two species. We classified GO and KEGG annotations of DEGs according to official classification and performed GO and pathway functional enrichment using topGO and phyper, both of which are functions from R. GO annotation and enrichment analysis was performed using an online-based software, DAVID.

## Ethics approval

Animal studies were approved by the University of Rochester Committee for animal resources.

## Consent for publication

Not applicable.

## Availability of data and materials

The datasets used and/or analyzed during the current study are available from the corresponding author on reasonable request.

## Author contributions

J.Z., X.T., X.F., and V.G. designed research. X.T., Z.Z., and E.R. generated stable cell lines and performed western blot and xenograft experiments. H.Z. constructed libraries and sequenced the samples. J.Z., Y.Z., Y.Y., B.R., A.C., X.S., and R.H. performed data control and bioinformatics analysis. E.N. provided critical samples. K.K. contributed to the result interpretation and paper writing. A.S., X.F., and V.G. supervised the study. X.T., X.F., and V.G. wrote the paper with input from all authors.

## Supplementary information

**Table S1.** Statistics of sequencing data and expressed genes.

| Sample | Number of Reads | Number of bases | Number of genes |
| --- | --- | --- | --- |
| BMR-1 | 34943168 | 5241475200 | 12816 |
| BMR-2 | 34237168 | 5135575200 | 12913 |
| BMR-3 | 34763768 | 5214565200 | 12783 |
| BMR-L-1 | 33980456 | 5097068400 | 12983 |
| BMR-L-2 | 34715316 | 5207297400 | 13185 |
| BMR-L-3 | 33697198 | 5054579700 | 13113 |
| BMR-L-R-1 | 34896340 | 5234451000 | 13370 |
| BMR-L-R-2 | 33554738 | 5033210700 | 13352 |
| BMR-L-R-3 | 35027026 | 5254053900 | 13428 |
| HCA2-1 | 34428058 | 5164208700 | 13549 |
| HCA2-2 | 34559592 | 5183938800 | 13533 |
| HCA2-3 | 34311928 | 5146789200 | 13570 |
| HCA2-T-1 | 34369286 | 5155392900 | 13537 |
| HCA2-T-2 | 34482134 | 5172320100 | 13531 |
| HCA2-T-3 | 34939600 | 5240940000 | 13546 |
| HCA2-T-LT-1 | 34654506 | 5198175900 | 13580 |
| HCA2-T-LT-2 | 34653804 | 5198070600 | 13517 |
| HCA2-T-LT-3 | 35401832 | 5310274800 | 13594 |
| HCA2-T-LT-RAS-1 | 34482018 | 5172302700 | 13755 |
| HCA2-T-LT-RAS-2 | 34917538 | 5237630700 | 13780 |
| HCA2-T-LT-RAS-3 | 35351198 | 5302679700 | 13771 |
| M-1 | 34856638 | 5228495700 | 13364 |
| M-2 | 33703474 | 5055521100 | 13165 |
| M-3 | 34872236 | 5230835400 | 13376 |
| M-L-1 | 34876306 | 5231445900 | 13423 |
| M-L-2 | 33989634 | 5098445100 | 13457 |
| M-L-3 | 34168260 | 5125239000 | 13452 |
| M-L-R-1 | 34267706 | 5140155900 | 13229 |
| M-L-R-2 | 33990568 | 5098585200 | 13355 |
| M-L-R-3 | 34491972 | 5173795800 | 13195 |
| NMR-1 | 34807890 | 5221183500 | 13144 |
| NMR-2 | 34095594 | 5114339100 | 13148 |
| NMR-3 | 34153410 | 5123011500 | 13339 |
| NMR-L-1 | 35874752 | 5381212800 | 13343 |
| NMR-L-2 | 36027014 | 5404052100 | 13436 |
| NMR-L-3 | 35718470 | 5357770500 | 13410 |
| NMR-L-R-1 | 35318112 | 5297716800 | 13443 |
| NMR-L-R-2 | 35524056 | 5328608400 | 13510 |
| NMR-L-R-3 | 34950234 | 5242535100 | 13508 |

**Table S2.** Number of orthologs between mouse and other species.

| BMR | NMR | Human | BMR+NMR | BMR+Human | NMR+Human | BMR+NMR+Human |
| --- | --- | --- | --- | --- | --- | --- |
| 15841 | 15838 | 16861 | 13907 | 14579 | 14870 | 13276 |

**Table S3.** Numbers of DEGs in LT+Ras transformed cells between species pairs.

| Mouse<br>VS<br>BMR | Mouse<br>VS<br>NMR | Mouse<br>VS<br>Human | BMR<br>VS<br>NMR | BMR<br>VS<br>Human | NMR<br>VS<br>Human |
| --- | --- | --- | --- | --- | --- |
| 2204 | 2515 | 2411 | 2440 | 2616 | 2459 |

**Figure S1.**
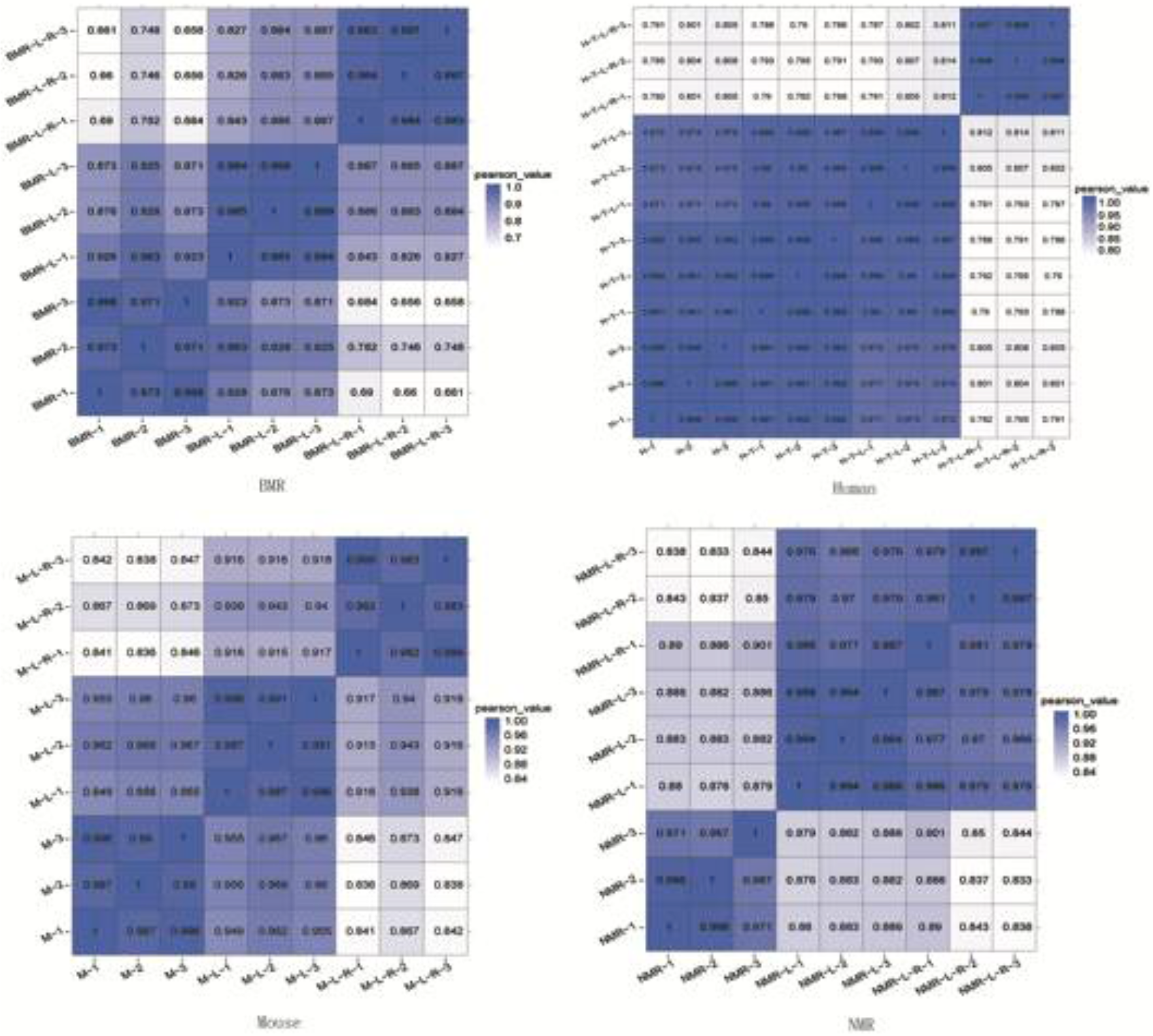
Pearson correlation coefficients between samples within each species. The heat maps were generated using the Pearson correlation coefficients for the RNA-seq data within each species.

**Figure S2.**
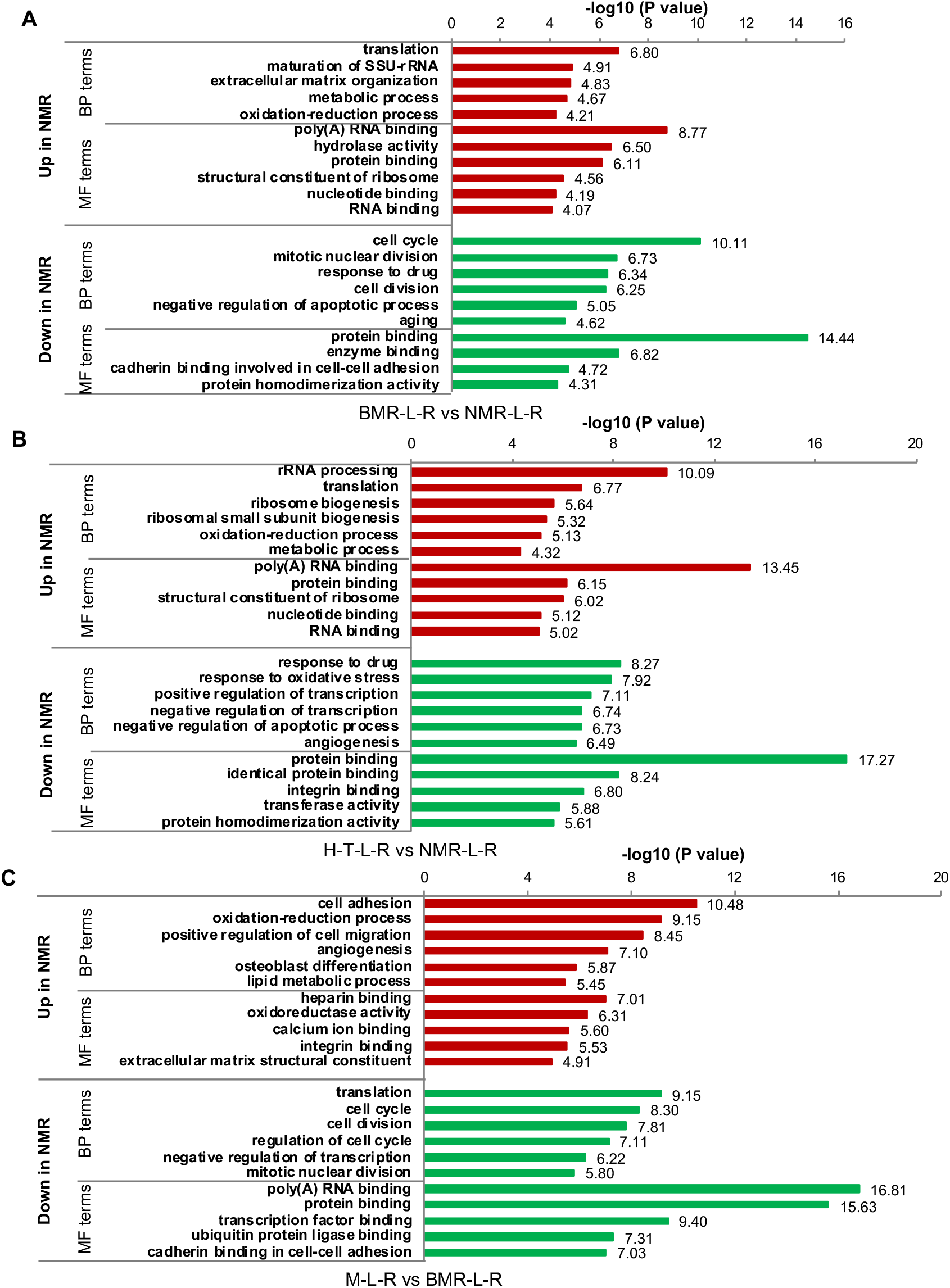
Gene expression variation in the LT and Ras-transformed cells across species. (**A**) GO enrichment analysis between the blind mole rat and the naked mole rat cells expressing Large T and H-Ras V12 (BMR-L-R vs NMR-L-R). (**B**) GO enrichment analysis between the human and the naked mole rat cells expressing Large T and H-Ras V12 (H-T-L-R vs NMR-L-R). (**C**) GO enrichment analysis between the mouse and the blind mole rat cells expressing Large T and H-Ras V12 (M-L-R vs BMR-L-R). Biological Process (BP) and Molecular Function (MF) terms with the most significant P values were plotted.

**Figure S3.**
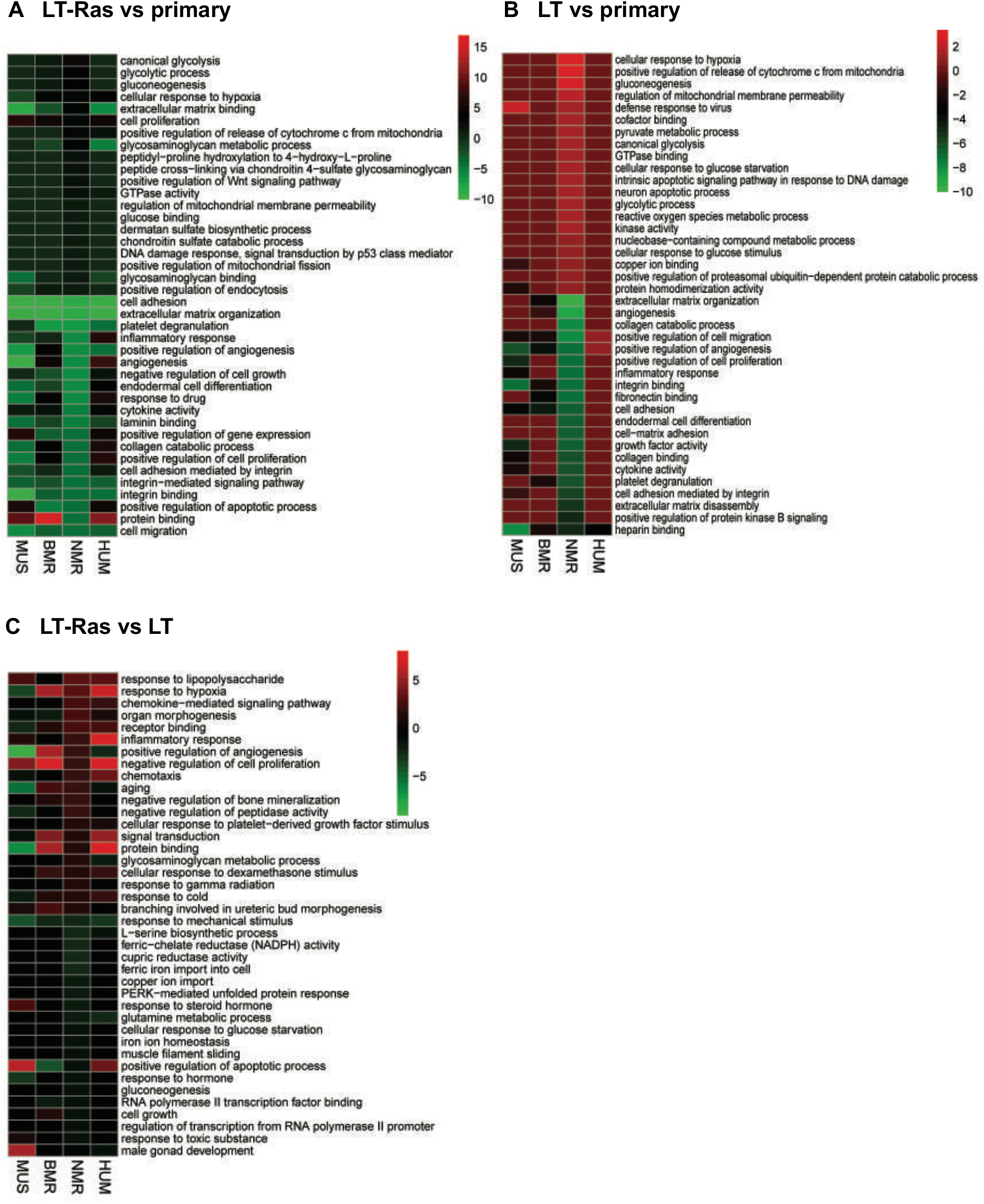
Differentially expressed genes between species induced by oncoprotein expression. (**A**) GO enrichment analysis of the DEGs induced by the combination of Large T (LT) and H-Ras V12 (Ras) across species. (**B**) GO enrichment analysis of the DEGs induced by LT across species. (**C**) GO enrichment analysis of the DEGs induced by H-Ras V12 across species. The same analyses as in Figure 3 were performed, with the GO terms arranged in the order of decreased significance in the NMR cells.

**Figure S4.**
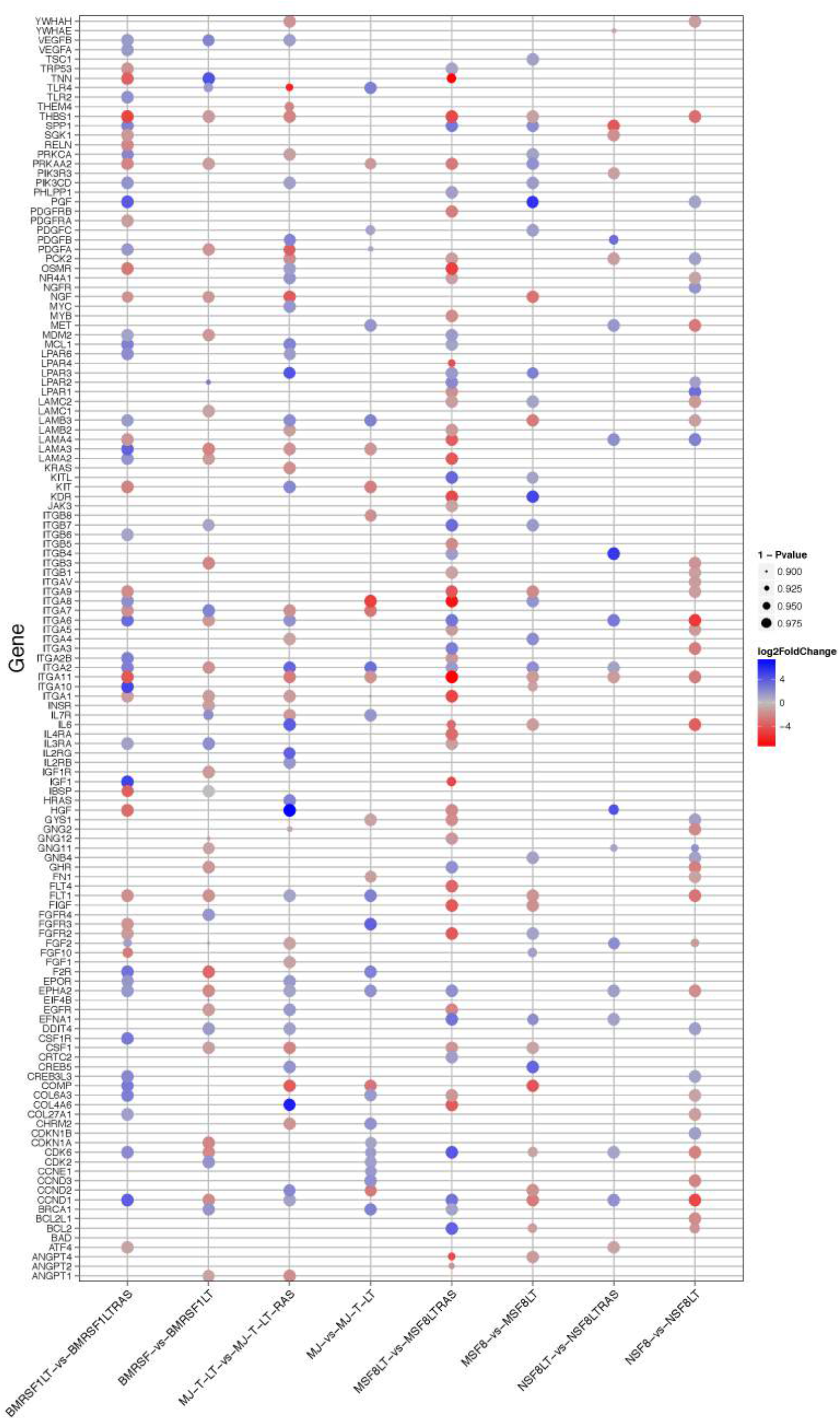
KEGG analysis of the PI3K pathway in oncogene-transformed cells. The genes of the PI3K pathway (KEGG: mmu04151) that show significant changes in the indicated sample pairs are shown. The size of the dots indicates the level of significance.

**Figure S5.**
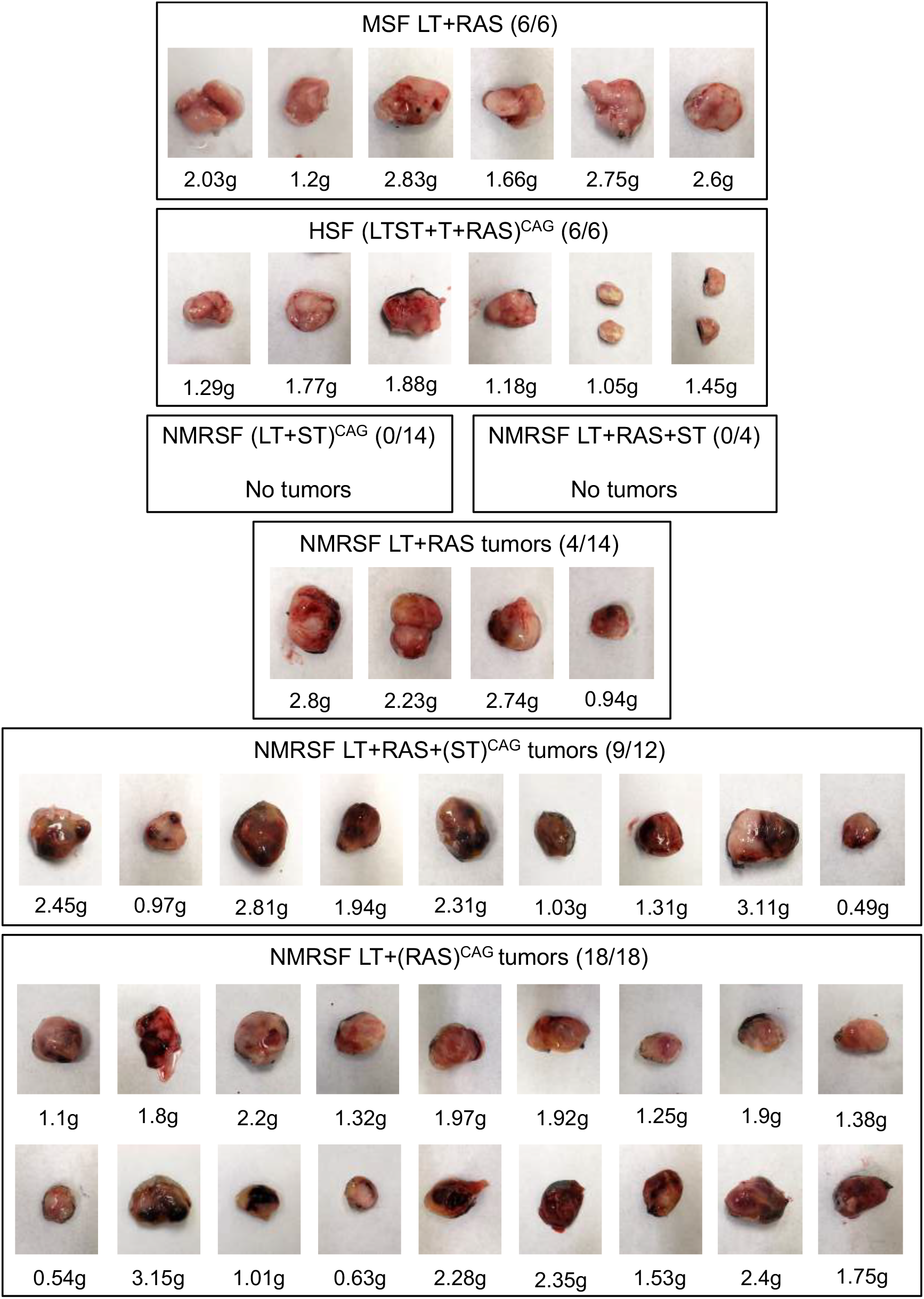
Tumors formed by injecting mouse, human and NMR cells expressing different combinations of oncoproteins in immunodeficient mice. Tumors formed from the xenograft experiments and their weights are shown. The numbers in parenthesis indicate number of tumors formed per the number of injection sites.

